# The TMCrys server for supporting crystallization of transmembrane proteins

**DOI:** 10.1101/446054

**Authors:** Julia K. Varga, Gábor E. Tusnády

**Affiliations:** “Momentum” Membrane Protein Bioinformatics Research Group, Institute of Enzymology, Research Center of Natural Sciences, Hungarian Academy of Sciences, H-1117 Budapest, Magyar tudósok körútja 2.

## Abstract

**Motivation:** Due to their special properties, the structures of transmembrane proteins are extremely hard to determine. Several methods exist to predict the propensity of successful completion of the structure determination process. However, available predictors incorporate data of any kind of proteins, hence they can hardly differentiate between crystallizable and non-crystallizable membrane proteins.

**Results:** We implemented a web server to simplify running TMCrys prediction method that was developed specifically to separate crystallizable and non-crystallizable proteins.

**Availability:** http://tmcrys.enzim.ttk.mta.hu

## 1 Introduction

Transmembrane proteins (TMP) play vital roles in the cells acting as gatekeepers and receptors in the cell and organelle membranes. They are frequently targeted by pharmaceuticals: a survey found that more than 50% of marketed drugs interact with TMPs (Hopkins and Groom, 2002). Although the human proteome consists of about 25% TMPs (Dobson *et al.,* 2015), however, of all known protein structures only 2% belong to them (Kozma *et al,* 2013) and less than a hundred human TMP non-redundant structure is determined (Varga *et al,* 2017). Knowing the structure of TMPs may aid drug development by providing targets for ligand screening and enabling the creation of models for proteins with unknown structures. However, membrane proteins reside in the cell membrane making the process of structure determination extremely difficult.

In the last 10 years, several prediction methods were developed to enhance the success of structure determination by estimating the chance of successful experiments. Most of them uses the data from TargetTrack (Berman *et al,* 2009) or its predecessors PepcDB and TargetDB (Chen *et al.,* 2004) and PDB structures (Kouranov *et al,* 2006). However, almost all of them mix globular and TM proteins leading to predict TMPs as ‘hard to crystallize’ (or somewhat equivalent) without the ability to distinguish between crystallizable and non-crystallizable TMPs. The only TMP-specific method is MEMEX (Martin-Galiano *et al.,* 2008) but being created in 2008, the data used is outdated. We introduced the TMCrys (Varga and Tusnády, 2018) method to aid the process of structure determination of TMPs. Since the algorithm of TMCrys requires installing some libraries and software packages hereby we introduce the TMCrys server, providing a graphical user interface for the prediction via our HPC to facilitate the usage of the method.

## 2 Methods

### 3.1 Introduction to TMCrys

Training and test datasets for TMCrys were created using PDBTM and TargetTrack databases as described in (Varga and Tusnády, 2018). Several physical and chemical features describing the sequences were calculated using the topology of the protein, predicted by CCTOP algorithm (Dobson *et al.,* 2015), and other programs (Xiao *et al.,* 2015; Overton and Barton, 2006; Petersen *et al.,* 2009; Walker, 2005). Three XGBoost Decision Trees models were trained to predict the success of purification, solubilization and crystallization, respectively. Finally, a model aggregating the results of the three steps were computed to predict the success of the whole process. The models were evaluated using 10-fold cross-validation and tested on their respective hold-out datasets.

### 3.2 Reliability of the predictions

Reliability of the prediction were defined as the distance from the threshold of the calculated probabilities, normalized to one:

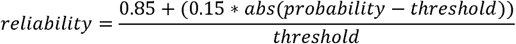

where threshold for the whole process was 0.85.

### 3.3 The TMCrys server

TMCrys server was developed using the Laravel web application framework (version 5.5.2) and designed with Bootstrap 3.2.7. Upon submitting a job, the sequences are forwarded to a highperformance computing (HPC) cluster. An Apache Axis server monitors the jobs on the cluster and provides the base of the communication between the HPC and the hosting server. The status of the job and the results are retrieved using SOAP requests. Several programs and scripts are run simultaneously to calculate features for the prediction to speed up the process. The results are sent back to the web server and displayed in HTML format and links are available for the download of the results in XML or tab separated format. Users may provide a job name for the identification of their job and optionally an email address as the results usually takes several minutes to obtain. An overview of the prediction process is provided in Supplementary Figure 1.

## 3 Results and discussion

### 3.4 Input

The server accepts input in several formats. Basically, one can submit sequences in FASTA format or space separated format. As the topology of the membrane protein is required for calculating the features, the user is permitted to submit topology of the protein calculated by themselves that should have the same length as the sequence and can contain the following labels: ‘I’ for inside, ‘M’ for membrane, ‘O’ for outside, ‘L’ for loops and ‘S’ for signal peptide (10). To avoid server overload, maximum 10 sequences can be submitted as one job. The sequences can also be uploaded in a single file.

### 3.5 Output

Three typical HTML outputs can be seen on Supplementary Figure 2. The server generates HTML output for all query proteins in the following format. A query protein appears in an expendable panel. The color of the panel gives information about the protein being membrane or non-membrane, the latter indicated with a yellow panel and ‘non-TMP’ label (Supplementary Figure 2C). When the protein was predicted to be membrane protein by CCTOP (or a topology was provided), a green or a red panel appears indicating whether the protein was predicted to be crystallizable (Supplementary Figure 2A) or non-crystallizable (Supplementary Figure 2B), respectively.

The predicted outputs are provided in numerical formats as well as a slider diagram, together with the reliability of the prediction. Besides the sequence and the topology of the query, similar entries from TargetTrack and TSTMP databases are also listed. The former ones aid the process by providing TargetTrack IDs of similar experiments already performed. The TSTMP is a database that collects human membrane proteins with existing structures that can be used for modeling the query protein (group 3D), membrane proteins that can be modeled (group Modelable) and proteins without existing structure or model (group ‘Target’). These latter proteins would become modelable if the structure of the query protein was solved. Last, some of the calculated features are also displayed, like instability index or average solvent accessible surface area.

The outputs can be downloaded in XML and tab-separated format, displaying all the above described features and outputs.

### 3.6 Direct interface

To enable programmatic access to TMCrys server a direct interface was established as well. The user can submit one sequence at a time with an ID and can monitor the progress of the job by calling a polling interface. The results can be downloaded in both tab or XML formats. A template script developed in Python, that can process multiprotein FASTA files, is also provided on the server.

## Acknowledgements

We thank László Dobson for creating the script for accessing the direct interface.

